# Temperature, photoperiod and life history traits in *Drosophila subobscura*

**DOI:** 10.1101/717967

**Authors:** Heidi J. MacLean, George W. Gilchrist

## Abstract

Temperature and photoperiod are generally reliable indicators of seasonality that have shaped the life histories of many temperate zone organisms. Anthropogenic climate change, however, may alter historical weather patterns and seasonal cues. Many studies have evaluated thermal effects on life history traits, but fewer have also examined photoperiodic effects. Because the degree of seasonal cue varies across latitude, we examine developmental plasticity in *Drosophila subobscura* populations sampled from latitudinal clines across Europe and North America. We examine the interaction between temperature and photoperiod on insect development time, adult survival, and fitness using a two by two factorial design with long (16L:8D) and short days (8L:16D) at high (23°C) and low temperatures (15°C).. We find that development time is dependent on both temperature and photoperiod but the low temperature/long day treatment revealed a dramatic and unexpected 4.5 day delay in eclosion. Fitness, estimated by the intrinsic rate of increase (r), showed a significant increase in response to temperature and a decrease in response to day length, and an interaction such that long-days reduced the effects of temperature. Additionally, cooler temperatures increased lifespan, and long-days reduced survivorship; temperature and day length interacted such that lifespan is relatively shorter in seasonally mismatched (long-cool, short-warm) conditions compared to matched conditions. These data highlight the importance of multiple abiotic factors in predicting species’ responses to climate change.

## Introduction

Temperature is a driving force in life history evolution, particularly for ectotherms (Roff, 1992, Tauber & Tauber, 1982). Temperature affects size, development time, lifespan, and fitness in insects (reviewed in Janisch, 1932, Dell et al., 2011, Angilletta, 2009). Although insects generally have a restricted range of body temperatures over which they can achieve high rates of growth, foraging and other aspects of fitness (Andrewartha & Birch, 1954, Magnuson et al., 1979, Huey & Hertz, 1984), they are nonetheless dispersed over a broad geographic range. Insects can adapt to local climatic conditions through heritable or developmentally plastic changes in morphology, voltinism, or physiology (Angiletta, 2009). Many populations also display reversibly plastic traits such as behavioral thermoregulation, diapause, and thermal hardening (Hoffmann, 2003, Kellermann et al., 2009, Mitchell et al., 2011). Both genetically and environmentally determined traits show predictable patterns with regard to climate along both latitudinal and elevation gradients (Watt, 1968, Gillis & Smeigh, 1987, Schultz et al., 1992, Berry & Willmer, 1986, Ellers & Boggs, 2004).

Temperature varies with some regularity along latitudinal gradients but photoperiod is a more reliable indicator of season in mid-to high latitude localities. Longer days are associated with summer and relatively warmer temperatures, while shorter days are associated with winter and relatively cooler temperatures. As a result, many organisms, including insects and plants, use photoperiod as a predictor of seasonal changes that can induce behavioral or physiological acclimation (Bradshaw & Lounibos, 1972, Marion, 1982, Paulsen, 1968). Photoperiod often cues dormancy or diapause, allowing organisms to conserve their energy and increase stress resistance during less hospitable months (Williams et al., 2014, Hoffmann & Sgrò, 2011, Tauber & Tauber, 1982). Both the duration and temperature during diapause can have direct consequences for life history (Williams et al., 2014, Bradshaw, 2004). As a result, most studies of photoperiodic effects on insects have focused on diapause (reviewed in Tauber & Tauber, 1982), with the exception of work in *Drosophila melanogaster* showing that long day photoperiod can decrease metabolic rate and increase thermal optima (Lanciani, 1990, Lanciani & Anderson, 1993, Lanciani et al., 1991). These researchers posit that the effects of photoperiod are adaptive, even compensatory, as they counteract the direct effects of temperature (Clarke, 2003, Lanciani et al., 1992).

Thermal compensation, or the ability of an organism to maintain physiological function in different thermal environments, appears to be induced by photoperiod and mediated through metabolic rates. Thermal compensation can allow for the conservation of energy in high temperature environments and increased performance in low temperature environments (Lanciani, 1990, Lanciani & Anderson, 1993, Lanciani et al., 1991). Although increased temperatures tend to decrease development time, decrease survival, and increase population growth rate, longer photoperiod can act to suppress metabolic rates. A study using damsel flies suggests that individuals reared in high temperature and long days develop more slowly than those in high temperature and short day environments (De Block & Stoks, 2003). If photoperiod is being used as a cue for compensatory modification of thermal sensitivity, then short days, might trigger a physiological decrease in development time, potentially through increased metabolic rate. Therefore, we hypothesize that at low temperatures, short days will decrease development time, decrease survival, and increase population growth rate relative to long days.

The broad seasonal and geographic range of many *Drosophila* species, coupled with well-developed experimental techniques and interesting ecological attributes makes members of this group compelling candidates to quantify any implications for life history traits. Here, we use *Drosophila subobscura* from the ancestral European and the colonizing North American populations (reviewed in Ayala et al., 1989a), sampled from three latitudes on two continents to quantify the effect of temperature, photoperiod, and their interaction on key life history traits. These populations exhibit clinal patterns in body size, wing length, and inversion polymorphism despite high rates of gene flow within continental populations (Balanya, 2006, Pascual et al., 2007, Huey et al., 2000, Gilchrist et al., 2001). The recent introduction of the North American population allows us to examine variation in plastic responses across latitudes and between continents. The theory of thermal compensation predicts that high latitude conditions are associated with reduced development times; this prediction is supported in common garden experiments (Śniegula et al., 2012). Latitudinal variation is of particular interest as recent anthropogenic climate change is projected to increased environmental stochasticity and temperature variability (Williams et al., 2007, IPCC, Climate Change 2007, Coumou & Rahmstorf, 2012), while photoperiod maintains its celestial regularity. Furthermore, global warming is associated with expansion of species ranges toward higher latitudes (Chen et al., 2011), moving populations into regions with more pronounced annual fluctuations in day length.

How does photoperiod affect the responses of life history and fitness traits in the context of the effects of temperature? To address this question, we reared flies from laboratory populations under four temperature-photoperiod regimes. We exposed replicated samples from six mass laboratory populations of *Drosophila subobscura* (three from each continent) to all combinations of temperature (High 23°C versus Low 15°C) and photoperiod (Long 16L:8D versus Short 16L:8D) treatments. The long photoperiod day is like a day in early May in northern Washington State, whereas the short photoperiod day is like a mid-December day at the same latitude. Thus, the factorial design created analogs of a warm summer day, a cool winter day, and the novel, mismatched conditions of a warm winter day, and a cool summer day. A control line was reared at standard laboratory conditions (18°C, 14 L:10 D). Eggs were reared to adulthood in each of these treatments and were monitored for length of development time. The adults were assayed for fitness (estimated by the intrinsic rate of increase, *r*) and survivorship. Temperature effects on drosophilids are well characterized: low temperatures are associated with longer developmental times, lower intrinsic rates of increase, and longer survivorship (Bateman, 1972, Hoffmann, 2003). However, many animals have the ability to adjust their physiology in anticipation of seasonal change. If photoperiodic effects on plasticity are interacting with temperature effects, then short days, which signal winter temperatures, might trigger seasonal thermal compensation thereby increasing metabolic rate and resulting in decreased development time, higher population growth rates and decreased survivorship. By quantifying the responses to different combinations of photoperiod and temperature, we will begin to tease apart the relative contributions of each variable in acclimation responses. The use of three populations from two continents allows me to examine geographic variability in these traits.

## Methods

### Study System

The ancestral populations of *D. subobscura* range from North Africa to Scandinavia, where they have been shaped by over 10,000 years of post-glacial environmental variability in their native habitat (Prevosti, 1955). In the late 1970’s *D. subobscura* colonized Puerto Monte, Chile (Brncic et al., 1981); genetic analysis suggests that the flies were Mediterranean in origin and suffered a severe bottleneck event (Pascual et al., 2007). Nonetheless, populations rapidly spread out from Puerto Monte, spanning over 10° of latitude by the mid 1980’s (Ayala et al., 1989b). *D. subobscura* were discovered on the western coast of North America in the early 1980’s (Beckenbach & Prevosti, 1986, Pascual et al., 2007), with populations ranging from Central California up the Pacific Coast and into Southern Canada. These flies appear to be derived from the introduced South American populations (Pascual et al., 2007). Despite the large geographic distance and the evidence of morphological clines, there is a high degree of gene flow among clinal populations of *D. subobscura* within each continent (Latorre et al., 1992, Pascual et al., 2001, Zivanovic et al., 2007). *D. subobscura* overwinter as adults and have no known diapause regardless of latitude of origin and thus no critical photoperiod (Lankinen, 1993).

The laboratory populations were established from approximately 20 gravid females collected from six natural populations along two parallel latitudinal clines from Europe (Aarhus, Denmark at 56.15°N, 10.22°E; Lille, France at 50.63°,N 3.07°E ; Valencia, Spain at 39.43°N, - 0.37°E) and North America (Port Hardy, British Colombia at 50.70°N, -127.42^a^E; Bellingham, Washington at 48.74°N, -122.47°E; and Gilroy, California at 37.01°N, -121.58°E) in the summer of 2004. They were maintained in the laboratory as large populations (N_e_>>200) at 18°C and 14L:10D cycle in population cages (25 cm X 14 cm X 12 cm) containing 100 ml of yeast/cornmeal/molasses media. Eggs were collected from these population cages within 18 hours of being laid. For each population, 500 eggs were distributed equally into 10 vials containing 10 ml of media. Two vials per population were then placed in each of the four experimental combinations of temperature (High 23°C versus Low 15°C) and photoperiod (Long 16L:8D versus Short 16L:8D) as well as the control condition (18°C and 14L:10D), and reared to adulthood. The flies were maintained in five Percival environmental chambers, each programmed for one of the five combinations of photoperiod and temperatures. Temperatures were monitored using Hobo data loggers within each chamber, with adjustments made to keep each chamber within ± 0.5°C of its target temperature throughout the daily cycle.

### Development Time

Development time was calculated from day of oviposition to the date and time of adult eclosion for each treatment. For each population, 500 eggs were distributed equally into 25 25×95mm vials with 10ml of media (20 eggs per vial). These vials were then distributed among the five photoperiod-temperature regimes and the control chamber. This was repeated over three days to ensure enough eggs and that the larvae were kept at low density to minimize competition. After pupation was observed, vials were checked every 8 hours (08:00, 16:00, 24:00) and any freshly emerged adults were counted and sexed. Each vial was scored until no more adults emerged.

### Survivorship

To quantify survivorship, 30 flies (15 male and 15 females) from each population and larval treatment were sexed during light CO_2_ anesthesia 48 hours after eclosion and placed in a survivorship cage and immediately returned to their rearing environment. A 25×95mm vial with 10ml of molasses-yeast agar medium with 0.15mg of active yeast was affixed to the side of the cage and replaced weekly. The flies were counted daily; dead individuals were removed and lost individuals were censored from the study. The study continued for 90 days and a censored survivorship model was used to account for the flies that remained alive or escaped in the course of the study.

### Intrinsic Rate of Increase

To quantify population growth rate, we employed the Model II serial transfer assay outlined in Muller and Ayala (1981). Flies from each of the six populations were reared at low density (20 eggs per 5mm vial) in each temperature and photoperiod combination (15°C, 8L:16D; 23°C, 8L:16D; 18°C 14L:10D; 15°C, 16L:8D; 23°C, 16L:8D). Adults were collected three to six days post-eclosion and sorted by sex into groups of 10, using light CO_2_ anesthesia. The flies were then returned to their experimental treatment conditions and given three days to recover before being combined with the opposite sex, for a total of 20 flies, in a 50 mL bottle of cornmeal-molasses agar with 23 mg of water diluted yeast paste. There were two vials per replicate and 3 replicates conducted from July 2008 through March 2009. Then, following the Model II serial transfer method (Mueller, 1981), all adults were removed, sexed, and counted once every seven days for eight weeks to obtain estimates the intrinsic rate of increase (*r)*.

For the Mueller assay, we first estimated lambda, the finite rate of increase or the rate at which a population size can change over one time step

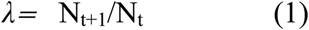

where N is the population size, and t is time (or generation). Population growth rate across experimental weeks is estimated by the linear equation

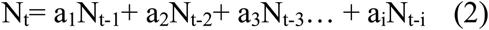

where a_i_ is the constant per capita output of an i-week old vial. As t gets larger, the per capita growth rate that is obtained is independent of the initial N and is estimated by the first positive eigenvalue of equation 2

We used a jackknife to estimate variance in λ between the replicates; we deleted the jth set of observations and calculated the leading eigenvalues (as above) yielding λ□_-j_. Then we calculated *m* pseudovalues as

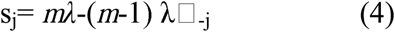

where j= 1,2,.. *m*.

The jackknifed estimate of the largest eigenvalue is the mean of the pseudovalues giving an estimate of λ□ for all replicates. (Mueller, 1981).

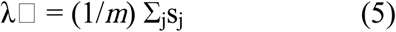

For overlapping generations over time, the intrinsic rate of increase (*r)* is estimated as:

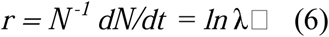

### Statistical Approach

All statistical analyses were performed in *R version 2*.*9*.*1* (Grambsch, 2014). For development time, we used a linear mixed effect model with the nlme package (Pinheiro, 2014) to assess the main and interactive effects of temperature, photoperiod, latitude and continent as predictor variables. This allowed us to estimate the main effects of temperature and photoperiod while testing for variability between continents and across latitudes. The predicted decrease in development time at higher temperatures would be indicated by a negative temperature term in the linear model. The predicted decrease in development time at short photoperiod would be indicated by significant positive effect of photoperiod in the linear model. If there is an interactive effect of temperature and photoperiod, we would expect to see an increase in development time on long days that is greater at colder temperatures. If there is clinal variation among populations, we would expect to find evidence of thermal compensation that is stronger with increasing latitude. Thus, we would expect that increasing latitude would shorten development time, and would be indicated by a negative latitude term in the linear model. Finally, if there is variation between continents, we would expect the continents to have different intercepts in the linear model.

We calculated and compared survival probabilities for each treatment using the “survreg” function in the “survival” library in *R version 2*.*9*.*1* (Grambsch, 2014). Day of death was examined as a function of population, temperature, and photoperiod. We tested for a main effect of continent and did not find one so it was used as a random effect or “strata” in the analysis. The predicted decrease in survival at higher temperatures would be indicated by a negative temperature term in the linear model. The predicted decrease in survival at short photoperiod would be indicated by significant positive effect of photoperiod in the linear model. If there is an interactive effect of temperature and photoperiod, we would expect to see an increase in survival on longer days that is greater at colder temperatures. If there is clinal variation among populations, we would expect to find evidence of thermal compensatory evolution, with compensation increasing with latitude. Thus, we would expect that increasing latitude would increase survival, and would be indicated by a positive latitude term in the linear model. Finally, if there is variation between continents, we would expect the continents to have different intercepts in the linear model.

For intrinsic rate of increase, we used the estimated *r* as the vial level response variable. We estimated *r* for all lines based on an eight week period starting with 20 adults. This was repeated three times for each line in each treatment. We again used a linear mixed effects model with the nlme package to assess the main and interactive effects of temperature, photoperiod and latitude and continent as predictor variables (Pinheiro, 2014). The predicted increase in population at higher temperatures would be indicated by a positive temperature term in the linear model. The predicted decrease in population growth rate at short photoperiod would be indicated by significant positive effect of photoperiod in the linear model. If there is an interactive effect of temperature and photoperiod, we would expect to see an increase in population growth rate on longer days that is greater at warmer temperatures. If there is clinal variation between populations, we would expect to find evidence of thermal compensation that is stronger with increasing latitude. Thus, we would expect that increasing latitude would increase growth rate, and would be indicated by a positive latitude term in the linear model. Finally, if there is variation between continents, we would expect the continents to have different intercepts in the linear model.

## Results

### Development Time

Development time in *D. subobscura* is longer than that of many other well-studied *Drosophila* species. The laboratory stock populations held at 14:10 LD and 18C typically take 20 days from egg to eclosion. Even though males eclose first in many holometabolic insects, we found no significant effect of sex on development time in any of the populations (figure 3, Sex: F_(1,60)_ =0.07, p =0.786). Nor did we observe an effect of continent, or latitude on development time (figure 3, Continent: F_(1,60)_ =0.35, p =0.55, Latitude: F_(1,60)_ =0.38, p =0.541). As expected, lower temperature (F_(1,60)_ =1279.09, p <<0.001) as well as long photoperiod (F_(1,60)_ =10.46, p =0.002) increased development time. The mean delay caused by longer photoperiod was only 0.34 days at high temperature but 4.79 days at low temperature, demonstrating the predicted interactive effect of temperature and photoperiod to delay development time (F_(1,60)_ =9.16, p =0.003) (Figure 1).

**Figure 1:**
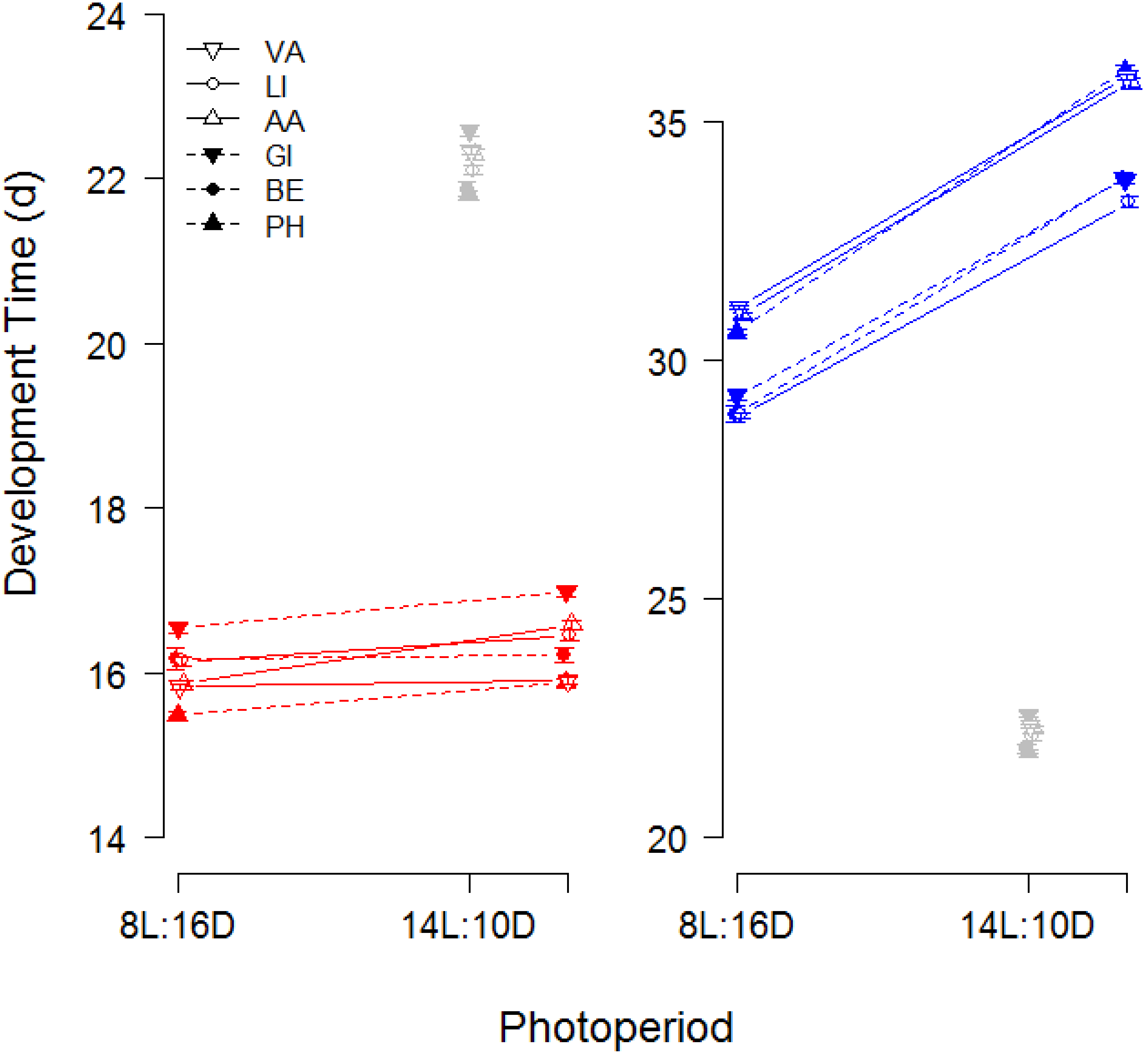
Development times in days with standard error bars. Temperature denoted in color-red being high temperature and blue being low temperature. Open circles (dashed lines) represent North American populations and closed circles (solid lines) represent European populations. Control treatment is shown in black.

**Figure 2:**
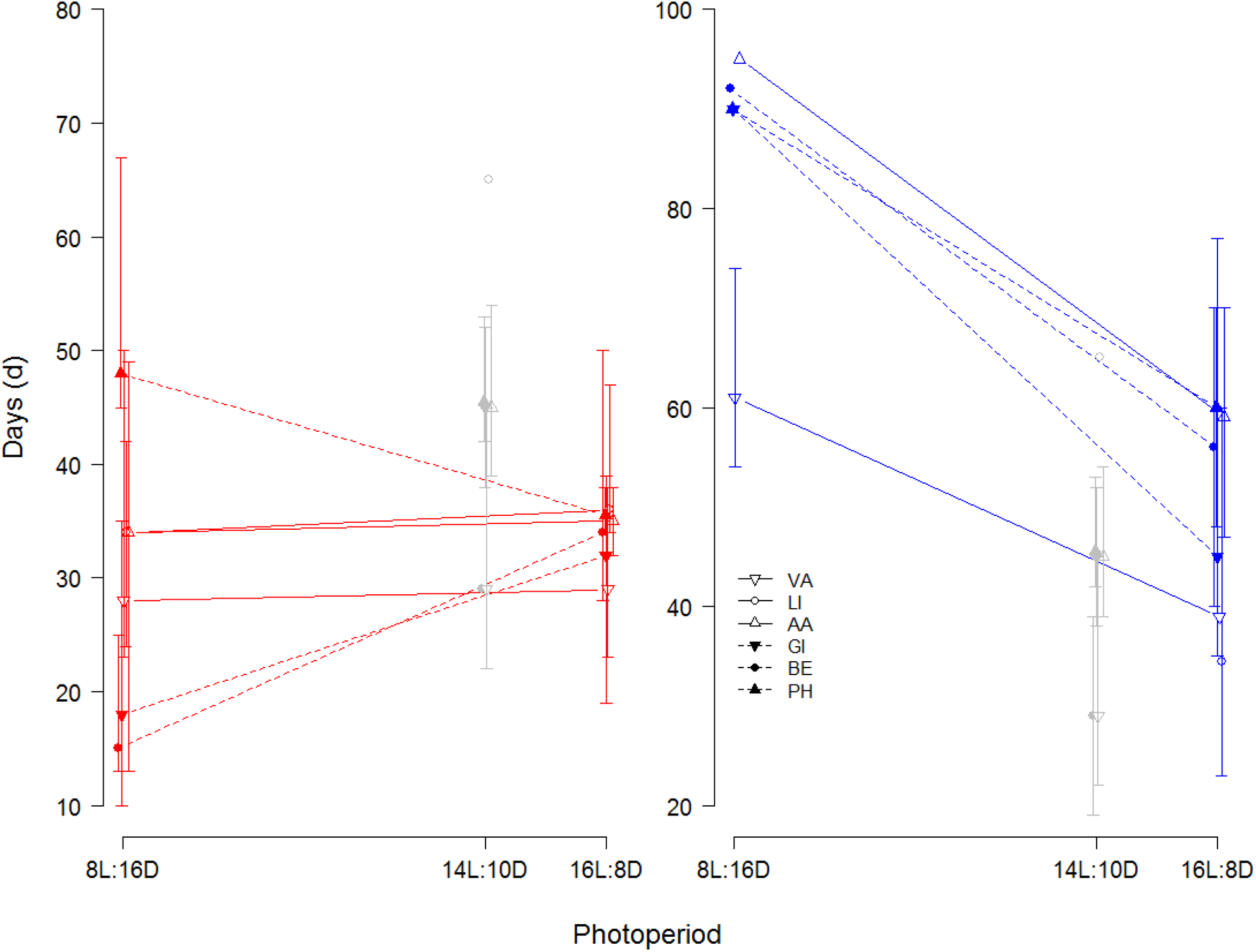
Average survival time of 50% of the population with 95% confidence intervals plotted as a function of photoperiod (denoted on the axis) and temperature (denoted in color-red being high temperature and blue being low temperature). Open circles (dashed lines) represent North American populations and closed circles (solid lines) represent European populations. Control treatment is shown in black

**Figure 3:**
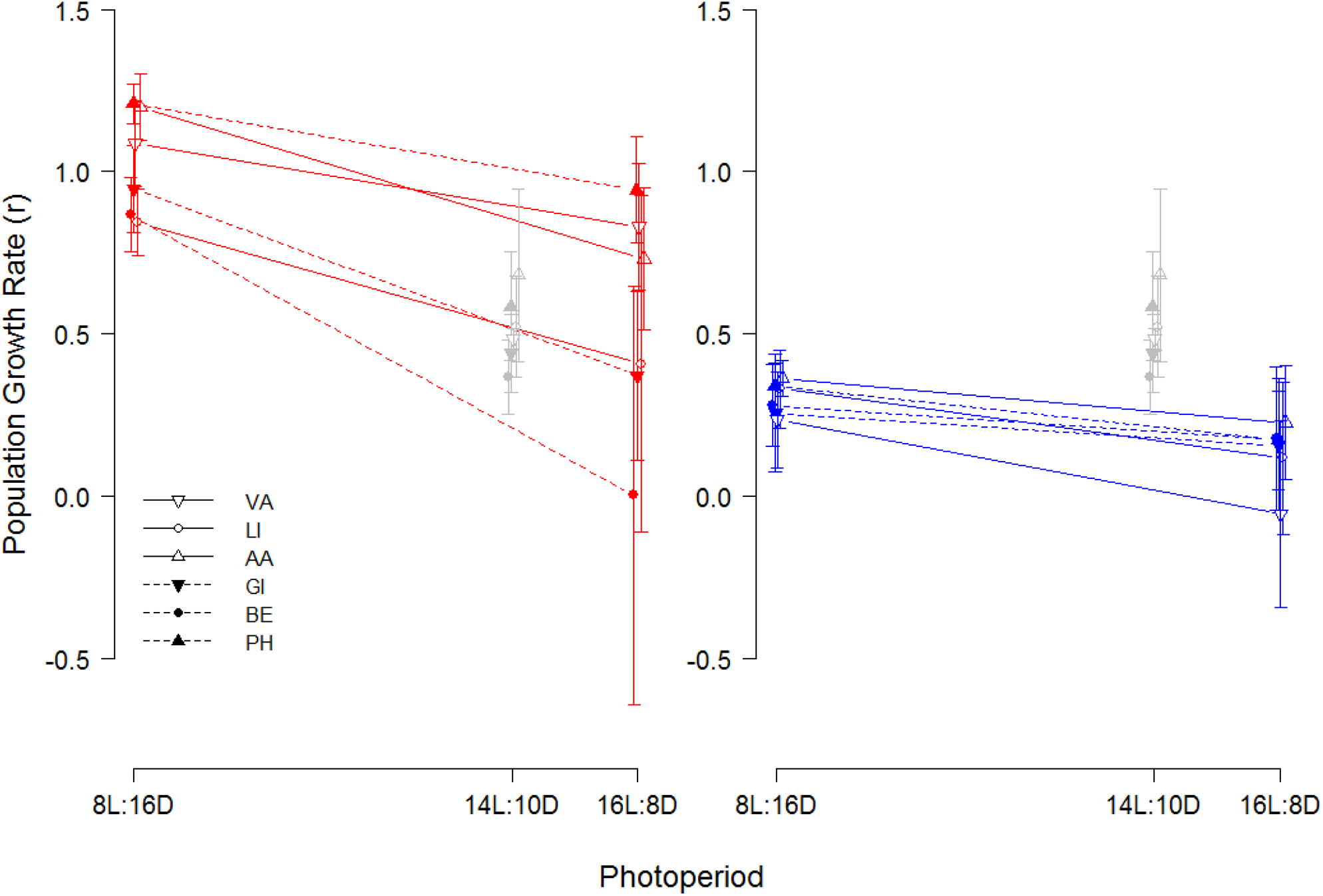
Intrinsic rate of increase with standard error bars plotted as a function of photoperiod (denoted on the axis) and temperature (denoted in color-red being high temperature and blue being low temperature). Open circles (dashed lines) represent North American populations and closed circles (solid lines) represent European populations. Control treatment is shown in black.

### Survivorship

While we found no significant effect of continent (deviance=1.30, df=1, p= 0.253); we did observe a positive effect of latitude on survival probabilities such that populations from higher latitudes survive longer on average (deviance=9.44, df=1, p = 0.002). As expected, survival is prolonged under both lower temperature (deviance=266.22, df=1, p <<0.001) and shorter days (deviance=47.80, df=1, p <<0.001). The interaction between temperature and photoperiod decreased survival in the high temperature and short day treatment (deviance=66.55, df=1, p <<0.001); however, contrary to our prediction, the low temperature and long day treatment showed lower survival relative to the low temperature and short day (Figure 2). In summary, we observed that the seasonally matched low temperature-short day and high temperature-long day treatments showed increased survivorship relative to the seasonally mismatched treatments.

### Intrinsic Rate of Increase

Our estimate of *r* did not vary with latitude (F_(5,144)_ =0.82, p= 0.537). As predicted, we observed a significant increase in population growth rate (r) with high temperatures (F_(1,144)_ =38.61 p<0.0001). However, population growth rate (r) decreased under long days (F_(1,144)_ =69.55, p<0.0001). Moreover, we saw a negative interaction between temperature and photoperiod such that high temperatures and short days produced the highest growth rate, and cold temperatures and long days produced the lowest growth rate (F_(1,144)_ =5.005, p = 0.0271. Figure 3).

## Discussion

Temperature is widely known to influence life history in insects in predictable ways (Hoffmann…), as was confirmed in this study. Hotter temperatures decreased *D. subobscura* development times and elevated the intrinsic rate of increase, but also decreased adult survival time. In our experimental treatment, the larvae in high temperature developed 11 to 15 days faster than those in low temperature, depending on photoperiod, regardless of latitude or continent of origin. Our measure of intrinsic rate of increase integrates both development time and reproductive output over an eight week period; high temperatures are associated with higher overall fitness. Previous research on *D. melanogaster* found that warmer temperatures increase reproductive output and decrease lifespan (Nunney & Cheung, 1997), as was observed in the present study.

Photoperiod is an important indicator of seasonality in the temperate zone; however, the effects of photoperiod on non-diapausing insects are little understood (Sheeba et al., 2001, Lanciani, 1990). Here we observe that longer photoperiod increased development time, shortened adult lifespan, and lowered the population growth rates; long days had a detrimental effect on all of our measured life history traits. One of the few prior studies of photoperiodic effects on traits other than diapause showed that for *D. melanogaster*, photoperiod can affect metabolic rates. Specifically, short photoperiods of 8L:16D elevate metabolic rates whereas long photoperiods of 16L:8D can suppress metabolic rates, which suggests thermal compensation (Lanciani, 1990, Lanciani et al., 1992, Clarke, 2003). Thermal compensation is generally adaptive and enables organisms to maintain their physiological function in the face of challenging thermal environments.

However, when we mismatched seasonal photoperiods and temperature, the results were not entirely as expected. Longer photoperiod significantly increased larval development time over short photoperiod at both test temperatures. Larval mass is the result of the acquisition and assimilation of resources; lower metabolic rate would increase the amount of time at each instar and delay overall development. We observed this developmental delay in both the development time experiment and in the intrinsic rate of increase experiment. Assuming that long photoperiods suppress metabolic rates, it would be as if the animals are consuming a poor quality diet; no matter how much they eat, they cannot metabolize it as efficiently as animals on shorter photoperiods. Diet quality manipulations in other insects, specifically *Manduca sexta*, demonstrate that lower quality diets significantly delay development (Davidowitz et al., 2004). Alternatively, if could be that feeding rates are slowed under longer photoperiods which could also lead to delayed development but further investigation is required to determine which mechanism is causing the delay.

Contrary to what we expected, long days were detrimental to survivorship at both temperatures in this study. Metabolic theory suggest that a lower metabolic rate contributes to a longer lifespan (reviewed in Finch, 1990). If long days decrease metabolic rate and if decreased metabolic rate increases life span, the prediction would be that longer days increase lifespan. This is completely unsupported by our data. Light increases activity in adult *D. melanogaster* (Martin et al., 1999), so longer days may increase the amount of active time and thus lead to increased resource use that is unlikely to be compensated by adult feeding behavior; this could reduce adult life span relative to the short day treatments. The negative effect on lifespan would be accentuated a high temperatures, which impose an additional kinetic energy cost. That cost could be reduced by compensation such that long days, which normally co-occur with higher temperatures, might reduce metabolic rate. In this experiment, that compensation was relatively ineffective; long days, even at low temperatures, reduced survival over short days.

Despite summer conditions generally promoting growth and development rates, population growth rate increased on shorter photoperiod at higher temperature. Given that shorter photoperiod decreased development time at both temperatures, we predicted that the intrinsic rate of increase might increase with higher temperature and shorter photoperiod. However, the observed decrease in development time was only a matter of hours at the high temperatures (Figure 1). This suggests that even though light increases activity in adult *D. melanogaster*, the increased metabolic rate of the adults may be associated with increased egg-laying and copulation behaviors regardless of the shorter lights-on time (Martin et al., 1999).

While we observed no continental differences in life history under these regimes, we did observe a clinal difference in survival, which is consistent with some degree of local adaptation in lifespan response to lower temperatures of the high latitude populations. Estimates of gene flow among the European populations indicate panmixia in nuclear DNA. In contrast, mitochondrial DNA has relatively low gene flow (Latorre et al., 1992), suggesting that males may disperse longer distances than females. While this could account for the lack evidence of local adaptation between latitudinal clines in plasticity for development time and population growth, it is more likely a lack of resolution in the methods used here. Moreover, we found no significant difference in life history responses to temperature and photoperiod cues between continents, suggesting that all of these populations may respond similarly to environmental cues.

This study is one of the first to explore the interaction between two of the most important environmental cues, photoperiod and temperature, in temperate zone clinal populations from two continents. Prior studies indicate that the high temperature treatments would probably shorten developmental times, increase population growth rates, and decrease survivorship (Bateman, 1972, Hoffmann, 2003), however little is known about the effects of photoperiod aside from some evidence of thermal compensation (Lanciani, 1990, Lanciani et al., 1992). Based on these studies, we expected short day photoperiod, which increases metabolic rates, to decrease development time, decrease survivorship, and increase population growth rates. In line with our prediction for development time, we observed that at 23°C, photoperiod has little effect, whereas at 15°C, the long days prolong development by three to five days. In contrast to our expectation that short days would increase survivorship, the interaction between temperature and photoperiod led to decreased survivorship for both of the mismatched conditions (short-warm, long-cool) relative to their matched counterparts. Perhaps the metabolic demands in mismatched conditions resulted in decreased survivorship, given Lanciani’s (1990) finding that a shorter photoperiod causes adaptive changes to counter the slowing kinetic effect of low temperature on metabolic rate (Conover & Present, 1990, Somero, 2004, Yamahira et al., 2007). Thus local adaptation in metabolic rate may account for the latitudinal differences observed in survival. Finally, in contrast with our prediction that short days would lead to a decrease in population growth rate at high temperatures, the greatest population growth rate (*r*) observed in this study was on short day photoperiod at 23°C.

These data introduce a new dimension to studies of thermal physiology. The quantification of thermal performance has generally been measured as a function of temperature and then extrapolated to ecological scenarios. Life history measures, such as the ones in this study, have become increasingly important in biophysical models of performance (Sears et al., 2011, Buckley & Kingsolver, 2012, Diamond et al., 2012). Failing to take into account the effect of photoperiod, or assuming that it is independent of temperature, could lead to dramatically incorrect estimations of performance in ectotherms under novel climatic regimes. Climate change presents a new suit of challenges for organisms that rely on the environment to cue seasonal changes in fitness-related traits (Bradshaw & Holzapfel, 2006, Parmesan, 2006). Novel temperature and photoperiod combinations could occur as a result of extreme or atypical weather patterns, both of which are expected to increase by 2100 (IPCC, Climate Change 2007, Williams et al., 2007). Moreover, species may experience novel temperature-photoperiod conditions as they shift their ranges poleward to seek thermal refuge (Chen et al., 2011, Crozier & Dwyer, 2006, Gorman et al., 2010, Wilson et al., 2005). As a result, some species are now experiencing longer summer and shorter winter photoperiods than ever before. These data suggest that simply considering temperature in calculating life history responses in these novel environments may not be sufficient. Organisms exist in a multivariate environment and modeling potential responses to climate change require integrating environmental information and not just looking at temperature to make predictions.

## Acknowledgements

We would like to thank John Swaddle, Joel Kingsolver, and Lauren Buckley for helpful comments on this manuscript. This work was supported by NSF grant DEB 0344273 to GWG.

